# Bacterial metabolism of tryptophan causes toxicity in *Caenorhabditis elegans* that is alleviated by sugar supplementation

**DOI:** 10.1101/2025.06.09.658597

**Authors:** Shivani Gahlot, Subodh, Jogender Singh

## Abstract

Tryptophan is an essential amino acid required not only for protein biosynthesis but also for the production of several physiologically important metabolites, including serotonin, kynurenine, and nicotinamide. Although dietary tryptophan is associated with various health benefits, excessive intake can result in adverse physiological effects. The specific tryptophan-derived metabolites responsible for such toxicity, however, remain incompletely characterized. Here, we investigate the mechanisms underlying tryptophan-induced toxicity in *Caenorhabditis elegans*. We observe that tryptophan concentrations of 1 mM or higher are highly toxic to *C. elegans*, blocking egg hatching. Notably, supplementation with various sugars, including glucose, fructose, mannose, galactose, rhamnose, and lactose, alleviates this toxicity. Genetic analyses reveal that host tryptophan metabolism is dispensable for the observed effects. Instead, bacterial metabolism, particularly the conversion of tryptophan to indole, is essential for mediating toxicity. Bacterial strains deficient in indole production abolished tryptophan-induced toxicity, and all sugars that conferred protection also suppressed bacterial indole synthesis. These findings demonstrate that tryptophan toxicity in *C. elegans* is primarily mediated by bacterial metabolism.

## Introduction

Tryptophan is an essential amino acid that is obtained exclusively through the diet. Beyond its fundamental role in protein biosynthesis, tryptophan regulates diverse physiological processes. It serves as a precursor for several bioactive molecules, including nicotinamide, serotonin, tryptamine, melatonin, and kynurenine, which participate in key metabolic and signaling pathways [1,2]. As the sole precursor of the neurotransmitter serotonin, tryptophan is vital for maintaining optimal neurological and physiological function [3]. Systemic or oral administration of tryptophan enhances serotonin synthesis [4], supporting its use as a dietary supplement to improve sleep quality and mood, and to manage psychiatric conditions such as depression and anxiety [5].

Emerging evidence suggests that dietary tryptophan and its metabolites may have therapeutic potential in various pathological conditions, including microbial infections, multiple sclerosis, inflammatory bowel disease, chronic kidney disease, and cardiovascular disease [2,4,6]. In rodent models, tryptophan supplementation has been shown to lower blood glucose levels in type 2 diabetic rats [7]. In *Caenorhabditis elegans*, elevated tryptophan levels, either through direct supplementation or inhibition of its catabolism, have been linked to increased lifespan [8–10]. Additional studies suggest that high dietary intake of tryptophan may protect against intestinal inflammation [11]. Tryptophan fortification of foods has also been associated with improved sleep in infants with sleep disorders, weight gain in malnourished children, and enhanced mood and sleep quality in elderly individuals [12,13].

Despite these benefits, excessive tryptophan intake has also been associated with adverse outcomes [14,15]. In rodents, dietary restriction of tryptophan has been shown to extend lifespan and delay the onset of age-associated pathologies, including tumor development [16,17]. Interestingly, the non-steroidal anti-inflammatory drug ibuprofen extends lifespan in yeast, worms, and flies by inhibiting tryptophan uptake [18]. Although a minor fraction of dietary tryptophan is utilized for serotonin and tryptamine synthesis, over 95% is catabolized via the kynurenine pathway [2,19]. Elevated levels of kynurenine pathway metabolites have been implicated in autoimmune diseases, neurodegenerative disorders, and various cancers [20]. Additionally, gut microbiota metabolize tryptophan into indole and its derivatives, which can be further converted in the liver into indoxyl sulfate, a uremic toxin associated with oxidative stress, tissue injury, and the progression of chronic kidney disease [6,21]. Given that tryptophan has both beneficial and detrimental effects, it is critical to elucidate the mechanisms underlying its toxicity and to better understand the regulation of its metabolism in host-microbe interactions.

In this study, we investigated the mechanisms underlying tryptophan toxicity in *C. elegans*. We found that tryptophan was highly toxic at concentrations of 1 mM or greater, leading to a complete inhibition of egg hatching. Transcriptomic analysis of tryptophan- exposed worms revealed a strong induction of genes associated with xenobiotic detoxification, oxidoreductase activity, and sugar transport. Given the transcriptional upregulation of sugar transport genes, we examined whether sugars modulate tryptophan toxicity. Remarkably, supplementation with several sugars, including glucose, fructose, mannose, galactose, rhamnose, and lactose, effectively rescued the toxicity. Genetic analyses of *C. elegans* mutants defective in tryptophan metabolism, including *tdo-2(ve552)*, *tph- 1(mg280)*, and *amx-2(ok1235)*, indicated that host-mediated catabolism was not responsible for the observed effects. Instead, bacterial metabolism of tryptophan played a central role in mediating toxicity. Specifically, the bacterial enzyme tryptophanase (TnaA), which converts tryptophan into indole, pyruvate, and ammonia, was required for toxicity. Worms cultured on bacterial diets unable to produce indole, including *Escherichia coli ΔtnaA* or *Pseudomonas aeruginosa*, showed complete resistance to tryptophan-induced toxicity. Notably, all sugars that alleviated toxicity also suppressed bacterial indole production. Our findings reveal that tryptophan toxicity in *C. elegans* is driven by bacterial metabolism and demonstrate the critical role of the host-microbiota axis in shaping the physiological outcomes of dietary amino acid intake.

## Results

### Tryptophan is highly toxic to *C. elegans* and inhibits egg hatching

While rearing *C. elegans* on *E. coli* OP50-seeded nematode growth medium (NGM) plates supplemented with various amino acids, we observed that tryptophan exhibited marked toxicity, completely inhibiting egg hatching. For assays on tryptophan-supplemented plates, adult worms were transferred for approximately 2 hours to allow egg laying, after which the adults were removed to obtain synchronized eggs (Materials and Methods). Egg hatching was scored after 24 hours of incubation. Representative images were acquired after 96 hours, when hatched eggs in control conditions had developed into gravid adults. At concentrations of 2 mM or higher, tryptophan completely inhibited egg hatching (Fig. 1A, B). This observation is consistent with a recent study [22], which reported that tryptophan exhibited the highest toxicity among the 20 standard amino acids, significantly impairing *C. elegans* development. In our assays, toxic effects were evident at 1 mM tryptophan, where more than half of the eggs failed to hatch, and at 2 mM, no hatching occurred (Fig. 1A, B). Moreover, tryptophan exposure also caused severe toxicity in adult worms, resulting in paralysis (Fig. 1C).

**Fig. 1.**
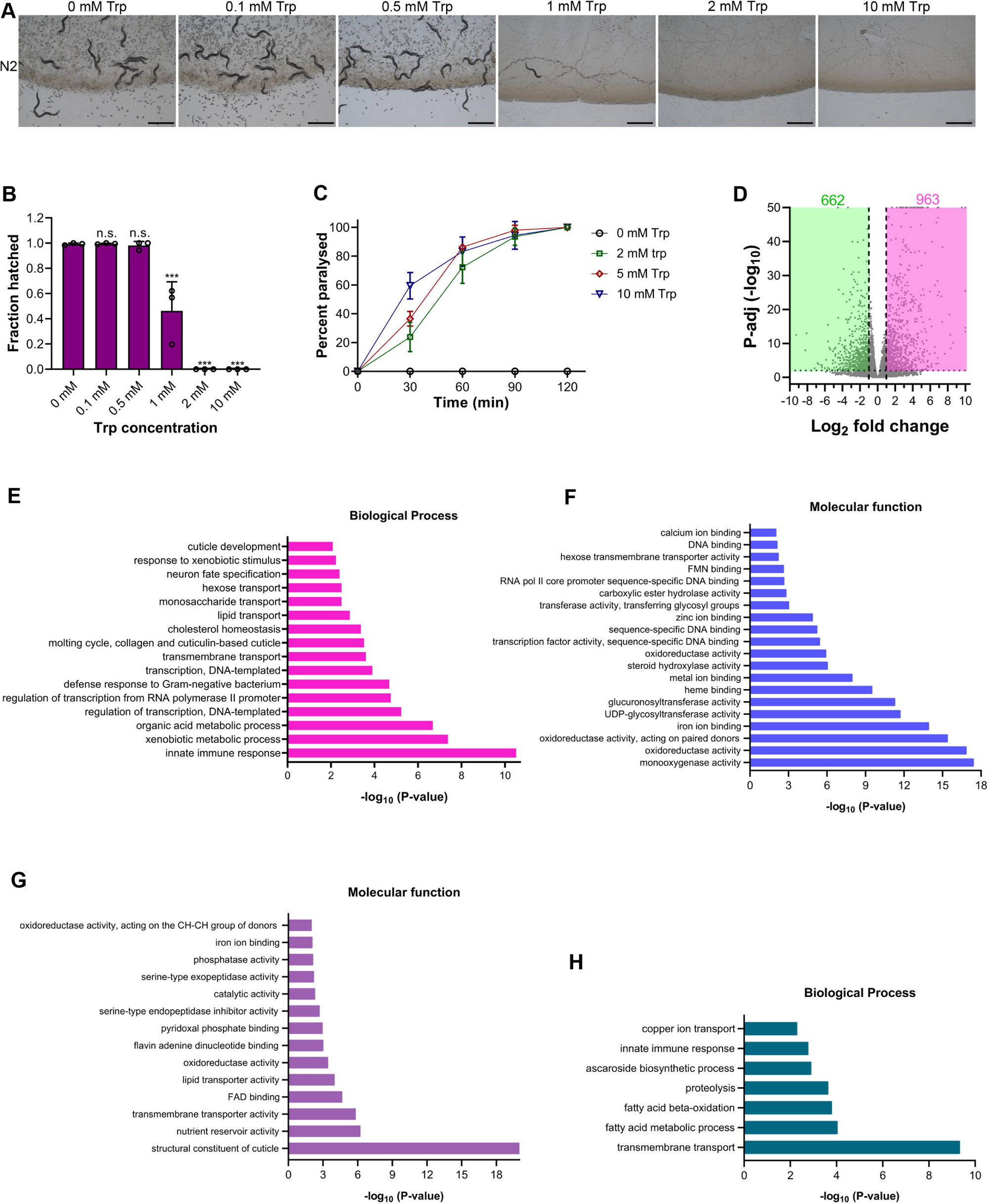
Tryptophan is highly toxic to *C. elegans* and inhibits egg hatching. (A) Representative images of N2 worms on various concentrations of tryptophan (Trp) on *E. coli* OP50 diet after 96 hours of hatching at 20°C. Scale bar = 1 mm. (B) Quantification of hatched eggs of N2 worms on various concentrations of tryptophan on *E. coli* OP50 after 24 hours of hatching at 20°C. \*\*\**P* < 0.001 and n.s., nonsignificant via ordinary one-way ANOVA followed by Dunnett’s multiple comparisons test (*n >*100 eggs for each condition from three independent biological replicates). (C) Quantification of paralysis of N2 worms on various concentrations of tryptophan on *E. coli* OP50 diet (*n >*150 adult worms for each condition from three independent biological replicates). (D) Volcano plot of upregulated and downregulated genes in N2 worms exposed to 2 mM tryptophan. (E)-(F) Gene ontology enrichment analysis of genes upregulated upon tryptophan exposure for biological process (E) and molecular function (F). (G)-(H) Gene ontology enrichment analysis of genes downregulated upon tryptophan exposure for molecular function (G) and biological process (H).

To investigate the mechanisms underlying tryptophan toxicity, we performed RNA sequencing to profile transcriptional changes following a 6-hour exposure of adult wild-type N2 worms to 2 mM tryptophan. Transcriptome analysis revealed 1,625 differentially expressed genes, including 963 upregulated and 662 downregulated transcripts (Fig. 1D, Table S1). Gene Ontology (GO) analysis of the upregulated genes revealed significant enrichment in biological processes related to innate immune and xenobiotic responses (Fig. 1E). In terms of molecular function, the upregulated genes were enriched for monooxygenase and oxidoreductase activities, including several members of the cytochrome P450 and UDP-glucuronosyltransferase (UGT) families (Fig. 1F). Notably, *ugt-34*, *ugt-33*, *cyp-35B2*, and *cyp-35A1* were among the most strongly induced genes. These findings suggested activation of xenobiotic detoxification pathways in response to tryptophan exposure. Conversely, GO analysis of downregulated genes indicated enrichment in terms associated with cuticle structure and transmembrane transport (Fig. 1G, H).

### Sugars rescue tryptophan toxicity

Interestingly, GO analysis also revealed that genes involved in monosaccharide and hexose transport were upregulated upon tryptophan exposure (Fig. 1E, F), suggesting a possible interaction between sugar metabolism and tryptophan toxicity. To explore this, we supplemented NGM plates containing 10 mM tryptophan with various monosaccharides at a concentration of 50 mM. Notably, while these monosaccharides alone did not affect *C. elegans* development in the absence of tryptophan (Fig. 2A), they fully rescued the toxic effects of tryptophan (Fig. 2B, C). Egg hatching was restored on plates containing 10 mM tryptophan supplemented with D-glucose, D-fructose, D-mannose, D-galactose, or L-rhamnose, whereas no hatching occurred on plates with 10 mM tryptophan alone.

**Fig. 2.**
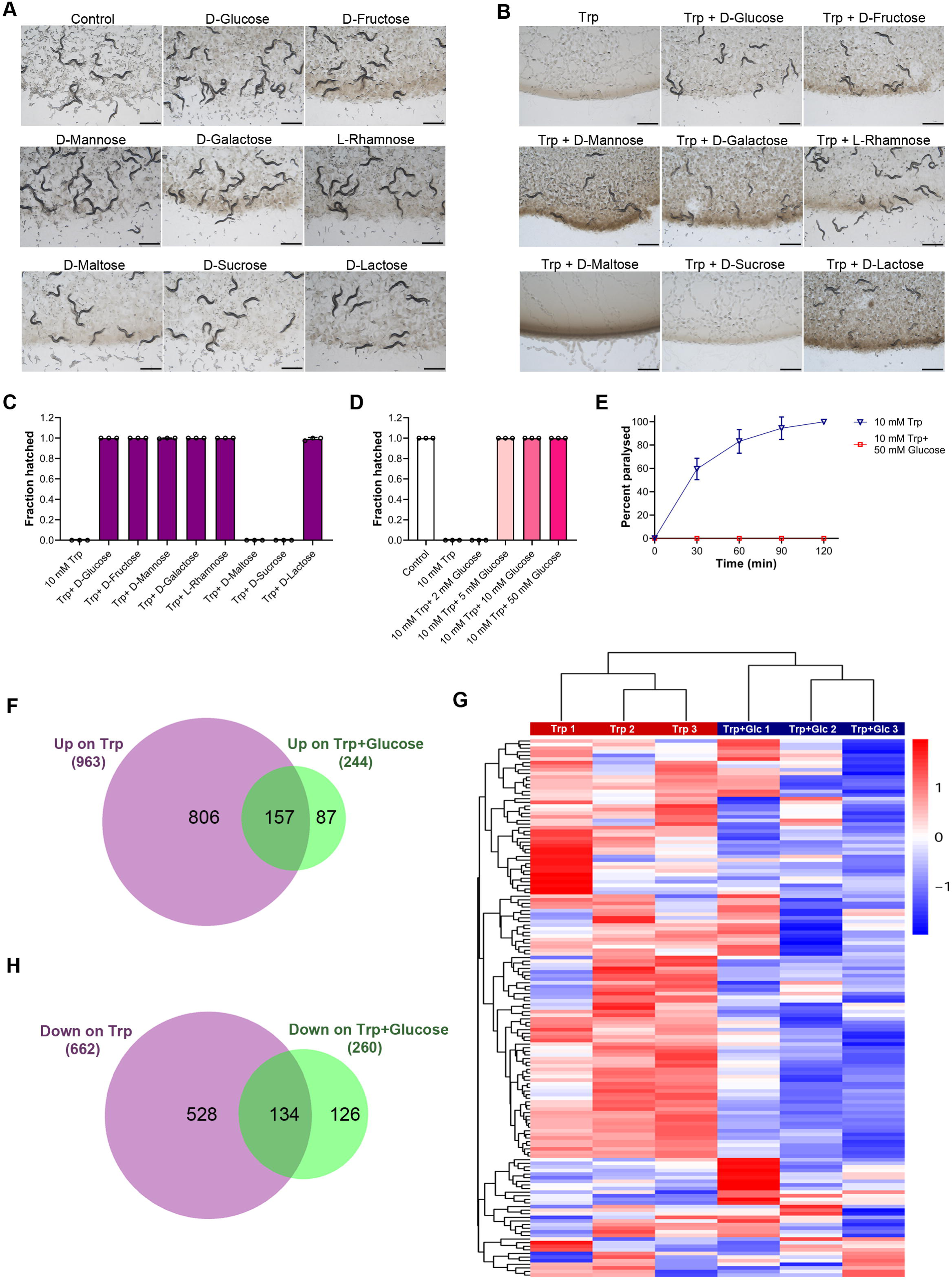
Sugars rescue tryptophan toxicity. (A) Representative images of N2 worms grown on *E. coli* OP50 diet for 96 hours at 20°C supplemented with 50 mM of different sugars. Scale bar = 1 mm. (B) Representative images of N2 worms grown on *E. coli* OP50 diet for 96 hours at 20°C containing 10 mM tryptophan (Trp) and 10 mM Trp supplemented with 50 mM of different sugars. Scale bar = 1 mm. (C) Quantification of hatched eggs of N2 worms grown on *E. coli* OP50 diet for 24 hours at 20°C containing 10 mM Trp and 10 mM Trp supplemented with 50 mM of different sugars (*n >*100 eggs for each condition from three independent biological replicates). (D) Quantification of hatched eggs of N2 worms grown on *E. coli* OP50 diet for 24 hours at 20°C containing 0 mM Trp (control), 10 mM Trp, and 10 mM Trp supplemented with different concentrations of glucose (*n >*150 eggs for each condition from three independent biological replicates). (E) Quantification of paralysis of N2 worms on 10 mM Trp and 10 mM Trp plus 50 mM glucose on *E. coli* OP50 diet (*n >*150 adult worms for each condition from three independent biological replicates). The 10 mM Trp data are the same as the data in Fig. 1C. (F) Venn diagram showing the overlap between the genes upregulated upon exposure to 2 mM Trp alone and 2 mM Trp plus 50 mM glucose. (G) Heat map showing the expression of upregulated genes common between Trp alone and Trp plus glucose (Trp+Glc). The fold change of 3 technical replicates of Trp and Trp+Glc with respect to the control is used for plotting. (H) Venn diagram showing the overlap between the genes downregulated upon exposure to 2 mM Trp alone and 2 mM Trp plus 50 mM glucose.

We further tested whether disaccharides could similarly suppress tryptophan toxicity. To this end, we supplemented tryptophan-containing plates with lactose, sucrose, or maltose. Interestingly, only lactose supplementation rescued tryptophan-induced toxicity (Fig. 2B, C). Neither sucrose nor maltose restored egg hatching in the presence of tryptophan. Importantly, all three disaccharides supported normal development in the absence of tryptophan (Fig. 2A), indicating that the inability of sucrose and maltose to rescue was not due to inherent developmental toxicity.

To determine whether lower concentrations of sugar could also rescue tryptophan toxicity, we supplemented 10 mM tryptophan plates with varying concentrations of glucose and assessed egg hatching. No rescue was observed at 2 mM glucose, whereas glucose concentrations of 5 mM and above fully restored egg hatching (Fig. 2D), demonstrating a dose-dependent protective effect. We further examined whether glucose supplementation could alleviate tryptophan toxicity in adult worms. Notably, the addition of 50 mM glucose completely rescued the paralysis phenotype observed in worms exposed to 10 mM tryptophan (Fig. 2E).

Next, we examined whether glucose supplementation could reverse the transcriptional changes induced by tryptophan exposure. RNA sequencing analysis of adult N2 worms exposed to 2 mM tryptophan plus 50 mM glucose for 6 hours showed a marked reduction in transcriptomic perturbation (Table S2). Only 244 genes were upregulated under the tryptophan-plus-glucose condition, compared to 963 genes upregulated in response to tryptophan alone (Fig. 2F). Notably, 806 of the genes upregulated in the tryptophan-only condition were no longer upregulated when glucose was present (Table S3). Moreover, most of the 157 genes upregulated in both conditions exhibited higher expression levels under tryptophan-only exposure than with tryptophan plus glucose (Fig. 2G). A similar trend was observed among downregulated genes, with a sharp reduction in the number of genes downregulated under the tryptophan-plus-glucose condition compared to tryptophan alone (Fig. 2H, Table S4). Collectively, these findings demonstrated that glucose supplementation effectively rescues tryptophan-induced toxicity at developmental, physiological, and transcriptional levels.

### Blocking tryptophan catabolism in *C. elegans* does not alleviate tryptophan toxicity

To understand how sugars mitigate tryptophan-induced toxicity, we first investigated whether excess tryptophan is metabolized into a toxic downstream product, and whether sugars act by suppressing the production or activity of this metabolite. In eukaryotes, tryptophan is primarily metabolized through the kynurenine and serotonin pathways (Fig. 3A). Although some eukaryotes, including *C. elegans*, can also generate indole from tryptophan [23], indole production is most prominently observed in bacteria [24]. The kynurenine pathway accounts for over 95% of dietary tryptophan metabolism in animals, and elevated kynurenine-to-tryptophan ratios have been associated with a variety of pathological conditions [2]. We therefore hypothesized that a toxic metabolite produced through the kynurenine pathway may underlie the observed developmental toxicity.

**Figure 3.**
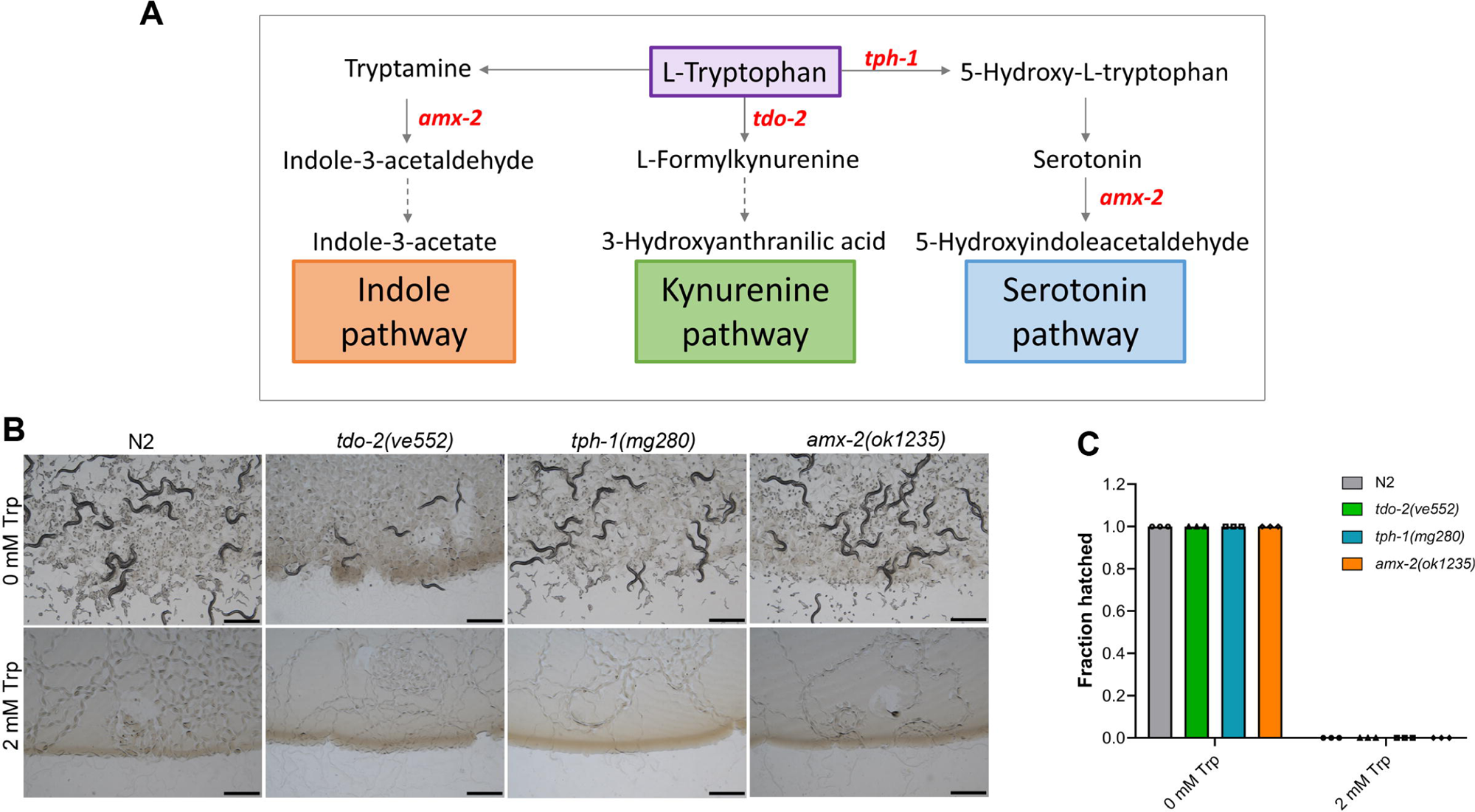
Blocking tryptophan catabolism in *C. elegans* does not alleviate tryptophan toxicity. (A) Simplified representation of tryptophan metabolism pathways. L-tryptophan is metabolized in *C. elegans* via three main pathways: the kynurenine, indole, and serotonin pathways. (B) Representative images of N2, *tdo-2(ve552)*, *tph-1(mg280)*, and *amx-2(ok1235)* worms after 96 hours of hatching at 20°C on *E. coli* OP50 diet containing 0 mM and 2 mM tryptophan (Trp). Scale bar = 1 mm. (C) Quantification of hatched eggs of N2, *tdo-2(ve552)*, *tph-1(mg280)*, and *amx-2(ok1235)* worms after 24 hours of hatching at 20°C on *E. coli* OP50 diet containing 0 mM and 2 mM Trp (*n >*90 eggs for each condition from three independent biological replicates).

To test this hypothesis, we examined *tdo-2* mutants, which lack tryptophan-2,3-dioxygenase, the rate-limiting enzyme that initiates kynurenine synthesis. If toxicity were mediated by kynurenine-derived metabolites, *tdo-2* loss-of-function mutants would be expected to exhibit resistance to tryptophan toxicity. However, *tdo-2(ve552)* mutants failed to hatch on 2 mM tryptophan, displaying sensitivity comparable to that of wild-type N2 animals (Fig. 3B, C).

We next asked whether the serotonin or indole branches of tryptophan metabolism contribute to toxicity. The enzyme TPH-1 catalyzes the first step in serotonin biosynthesis, whereas AMX-2 oxidizes amines such as serotonin and tryptamine to 5-hydroxyindole acetaldehyde and indole-3-acetaldehyde, respectively (Fig. 3A) [25,26]. Eggs from both *tph-1(mg280)* and *amx-2(ok1235)* mutants failed to hatch on 2 mM tryptophan, showing no increased resistance relative to N2 (Fig. 3B, C). Taken together, these results indicated that blocking host-mediated tryptophan catabolic pathways does not rescue tryptophan toxicity in *C. elegans*.

### Bacterial metabolism of tryptophan is essential for its toxicity in *C. elegans*

Since host-mediated tryptophan catabolism did not appear to underlie tryptophan toxicity, we investigated whether bacterial metabolism of tryptophan contributes to the toxic effects observed in *C. elegans*. To test this, we transferred *C. elegans* eggs to unseeded NGM plates containing 10 mM tryptophan and monitored their hatching. It is well established that *C. elegans* eggs can hatch in the absence of food and arrest at the L1 larval stage. While eggs fail to hatch on *E. coli* OP50-seeded plates containing 10 mM tryptophan, the use of unseeded plates enables assessment of tryptophan toxicity in the absence of bacterial metabolism.

Strikingly, all eggs transferred to unseeded plates containing 10 mM tryptophan hatched successfully (Fig. 4A, B). This observation strongly suggested that active bacterial metabolism is required for tryptophan-induced toxicity.

**Figure 4.**
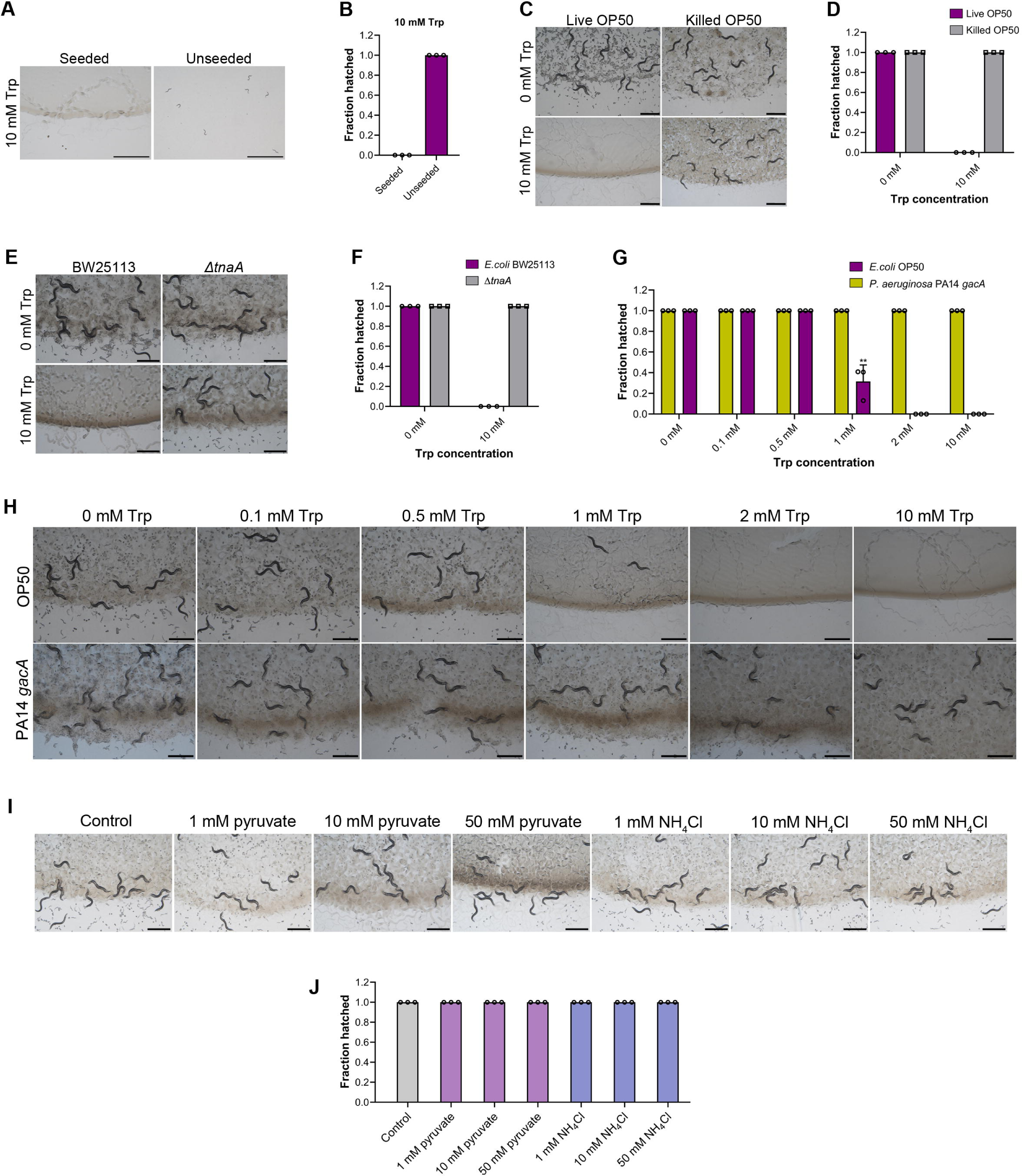
Bacterial metabolism of tryptophan is essential for its toxicity in *C. elegans*. (A) Representative images of N2 worms after 24 hours of hatching at 20°C on unseeded and *E. coli* OP50-seeded plates supplemented with 10 mM tryptophan (Trp). Scale bar = 1 mm. (B) Quantification of hatched eggs of N2 worms after 24 hours of hatching at 20°C on unseeded and *E. coli* OP50-seeded plates supplemented with 10 mM Trp (*n >*100 eggs for each condition from three independent biological replicates). (C) Representative images of N2 worms after 96 hours of hatching at 20°C on live and kanamycin-killed *E. coli* OP50 diet containing 0 mM and 10 mM Trp. Scale bar = 1 mm. (D) Quantification of hatched eggs of N2 worms after 24 hours of hatching at 20°C on live and kanamycin-killed *E. coli* OP50 diet containing 0 mM and 10 mM Trp (*n >*200 eggs for each condition from three independent biological replicates). (E) Representative images of N2 worms after 96 hours of hatching at 20°C on wild-type *E. coli* BW25113 and *ΔtnaA* mutant diet containing 0 mM and 10 mM Trp. Scale bar = 1 mm. (F) Quantification of hatched eggs of N2 worms after 24 hours of hatching at 20°C on wild- type *E. coli* BW25113 and *ΔtnaA* mutant diet containing 0 mM and 10 mM Trp (*n >*80 eggs for each condition from three independent biological replicates). (G) Quantification of hatched eggs of N2 worms after 24 hours of hatching at 20°C on various concentrations of Trp on *E. coli* OP50 and *P. aeruginosa* PA14 *gacA* mutant. \*\*\**P* < 0.001 via the *t*-test (*n >*100 eggs for each condition from three independent biological replicates). Because *P. aeruginosa* virulence might affect the hatching and development of *C. elegans*, we used the *gacA* mutant, which has drastically reduced virulence [73]. (H) Representative images of N2 worms after 96 hours of hatching at 20°C on various concentrations of Trp on *E. coli* OP50 and *P. aeruginosa* PA14 *gacA* mutant. Scale bar = 1 mm. (I) Representative images of N2 worms on various concentrations of sodium pyruvate and ammonium chloride on *E. coli* OP50 diet after 96 hours of hatching at 20°C. Scale bar = 1 mm. (J) Quantification of hatched eggs of N2 worms on various concentrations of sodium pyruvate and ammonium chloride on *E. coli* OP50 diet after 24 hours of hatching at 20°C (*n >*200 eggs for each condition from three independent biological replicates).

To validate this finding, we monitored the hatching of *C. elegans* eggs on 10 mM tryptophan plates seeded with kanamycin-killed *E. coli* OP50. Notably, in the presence of dead bacteria, tryptophan did not impair egg hatching or subsequent development; all worms developed to become fertile adults (Fig. 4C, D). Together, these results indicated that active bacterial metabolism of tryptophan generates a toxic product that mediates the observed effects in *C. elegans*.

In *E. coli*, tryptophan is primarily metabolized via the indole pathway [24].

Specifically, the bacterial enzyme TnaA catalyzes the conversion of tryptophan into indole, pyruvate, and ammonia [27]. We therefore hypothesized that *E. coli* TnaA-mediated metabolism of tryptophan is responsible for the observed toxicity. To test this, we utilized an *E. coli ΔtnaA* mutant, which lacks tryptophanase, and assessed *C. elegans* egg hatching on 10 mM tryptophan plates seeded with this mutant strain. Strikingly, all eggs hatched and developed normally on 10 mM tryptophan plates seeded with *E. coli ΔtnaA* (Fig. 4E, F), confirming that the toxic effect of tryptophan required its metabolism by bacterial TnaA.

To further substantiate this conclusion, we examined whether tryptophan was toxic when metabolized by a bacterium that does not produce tryptophanase. *P. aeruginosa* catabolizes tryptophan via the oxidative kynurenine pathway, generating anthranilate rather than indole [28]. When *C. elegans* eggs were cultured on 10 mM tryptophan plates seeded with *P. aeruginosa*, all eggs hatched, and no toxicity was observed (Fig. 4G, H).

Since TnaA catalyzes the conversion of tryptophan into indole, pyruvate, and ammonia, we next examined whether pyruvate or ammonia could independently elicit toxicity. Supplementation with up to 50 mM pyruvate or ammonia did not affect egg hatching, indicating that neither metabolite contributes to the observed toxicity (Fig. 4I, J). Together, these findings demonstrated that the metabolism of tryptophan by bacterial tryptophanase, most likely through the production of indole, is responsible for the toxic effects of tryptophan in *C. elegans*.

### Indole causes toxicity in *C. elegans*

To directly assess whether indole causes toxicity in *C. elegans*, we supplemented NGM plates with indole and examined its effect on egg hatching. Similar to the effects observed with tryptophan, indole supplementation on *E. coli* OP50-seeded plates inhibited egg hatching, with toxic effects evident at concentrations of 1 mM and above (Fig. 5A, B). Since *P. aeruginosa* rescued the toxic effects of tryptophan, we next investigated whether it could similarly attenuate indole-induced toxicity. However, indole remained toxic on *P. aeruginosa*- seeded plates, and no eggs hatched at concentrations of 2 mM or higher (Fig. 5A, B). At 1 mM indole, a significantly greater number of eggs hatched on *P. aeruginosa*-seeded plates compared to those seeded with *E. coli*. This difference is likely attributable to the endogenous production of indole by *E. coli*, which *P. aeruginosa* does not produce, resulting in a higher effective concentration of indole in the *E. coli*-seeded condition.

**Figure 5.**
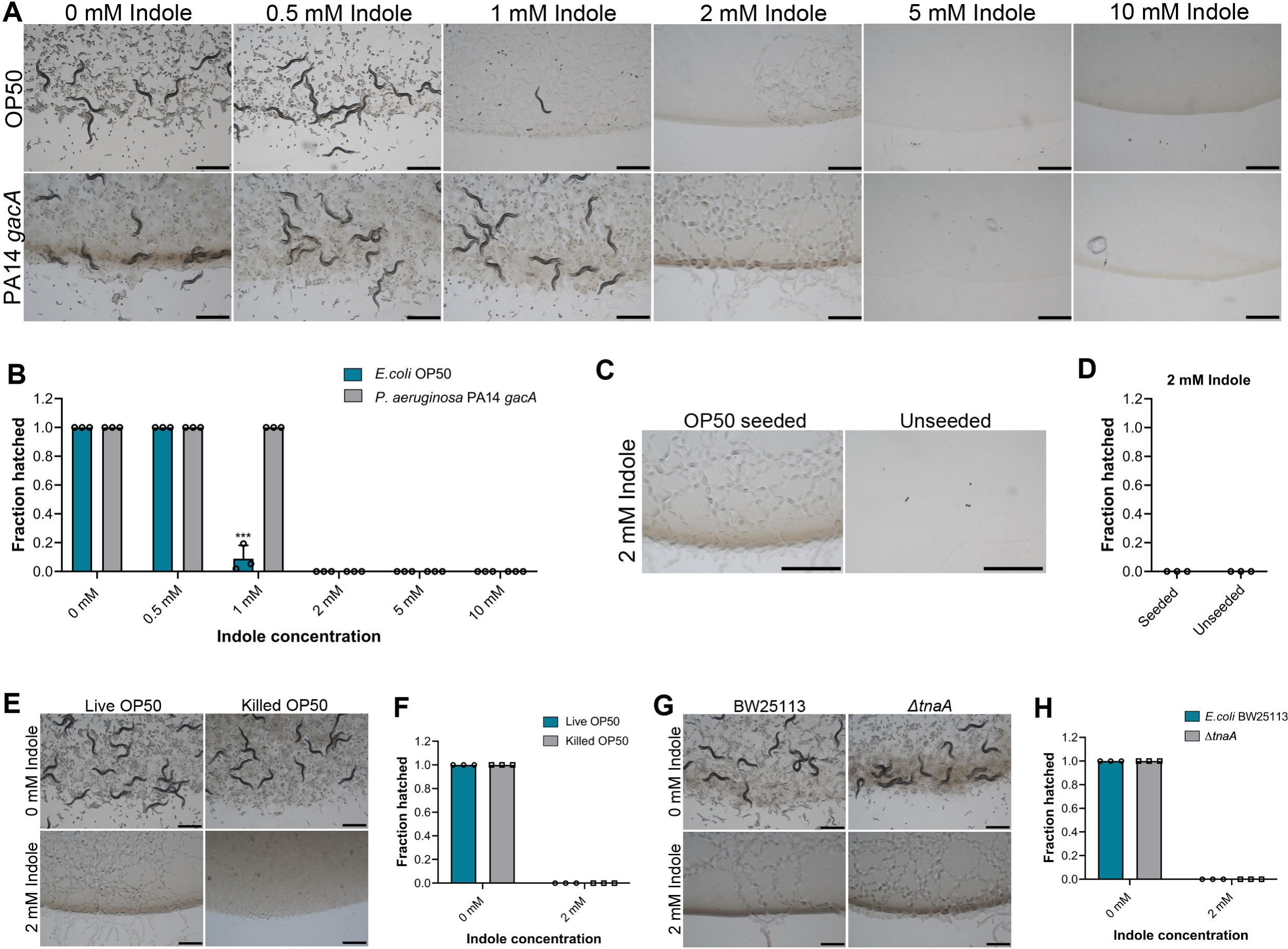
Indole causes toxicity in *C. elegans*. (A) Representative images of N2 worms after 96 hours of hatching at 20°C on various concentrations of indole on *E. coli* OP50 and *P. aeruginosa* PA14 *gacA* mutant. Scale bar = 1 mm. (B) Quantification of hatched eggs of N2 worms after 24 hours of hatching at 20°C on various concentrations of indole on *E. coli* OP50 and *P. aeruginosa* PA14 *gacA* mutant. \*\*\**P* < 0.001 via the *t*-test (*n >*70 eggs for each condition from three independent biological replicates). (C) Representative images of N2 worms after 24 hours of hatching at 20°C on unseeded and *E. coli* OP50-seeded plates supplemented with 2 mM indole. Scale bar = 1 mm. (D) Quantification of hatched eggs of N2 worms after 24 hours of hatching at 20°C on unseeded and *E. coli* OP50-seeded plates supplemented with 2 mM indole (*n >*40 eggs for each condition from three independent biological replicates). (E) Representative images of N2 worms after 96 hours of hatching at 20°C on live and kanamycin-killed *E. coli* OP50 diet containing 0 mM and 2 mM indole. Scale bar = 1 mm. (F) Quantification of hatched eggs of N2 worms after 24 hours of hatching at 20°C on live and kanamycin-killed *E. coli* OP50 diet containing 0 mM and 2 mM indole (*n >*200 eggs for each condition from three independent biological replicates). (G) Representative images of N2 worms after 96 hours of hatching at 20°C on wild-type *E. coli* BW25113 and *ΔtnaA* mutant diet containing 0 mM and 2 mM indole. Scale bar = 1 mm. (H) Quantification of hatched eggs of N2 worms after 24 hours of hatching at 20°C on wild- type *E. coli* BW25113 and *ΔtnaA* mutant diet containing 0 mM and 2 mM indole (*n >*200 eggs for each condition from three independent biological replicates).

To determine whether bacterial presence is required for indole toxicity, we examined egg hatching on both unseeded and *E. coli*-seeded indole plates. Notably, egg hatching was similarly inhibited on unseeded plates containing 2 mM indole (Fig. 5C, D), indicating that indole exerts its toxic effects independently of live bacteria. Similarly, indole inhibited egg hatching on plates seeded with kanamycin-killed *E. coli* (Fig. 5E, F), further confirming that bacterial viability is not required for indole-induced toxicity.

We also tested whether the *E. coli ΔtnaA* mutant, which lacks tryptophanase and cannot convert tryptophan to indole, could alter indole toxicity. Importantly, the *ΔtnaA* mutant did not mitigate indole toxicity, as eggs failed to hatch on 2 mM indole plates seeded with this strain (Fig. 5G, H). Collectively, these findings demonstrate that indole itself is inherently toxic to *C. elegans*, and this toxicity is independent of bacterial metabolism. While *E. coli*-derived indole mediates the toxic effects of tryptophan, the presence of indole alone is sufficient to exert toxicity, irrespective of bacterial species or metabolic activity.

### Indole derivatives cause toxicity in *C. elegans*

Experiments on unseeded plates demonstrated that indole, in the absence of bacterial metabolism, is intrinsically toxic to *C. elegans* and inhibits egg hatching. In the human gut, indole is further metabolized by bacteria into several derivatives [29]. To examine whether these human-relevant metabolites also exert toxic effects in *C. elegans*, we tested a panel of common indole derivatives.

Although none were as toxic as indole itself, all derivatives exhibited measurable toxicity (Fig. 6). Indole-3-acetic acid and indole-3-carboxaldehyde impaired egg hatching at concentrations of 5 mM and above (Fig. 6A). Indole-3-propionic acid reduced hatching at 10 mM, whereas indole-3-butyric acid showed no effect up to 10 mM. In addition to embryonic toxicity, all derivatives disrupted larval development. Notably, no larvae reached adulthood on 5 mM indole-3-carboxaldehyde, and all derivatives completely inhibited development at 10 mM (Fig. 6B). Collectively, these findings demonstrated that indole derivatives relevant to human metabolism are also toxic to *C. elegans*, affecting both embryonic and post-embryonic development. Their reduced potency relative to indole likely reflects decreased membrane permeability due to the presence of polar functional groups [30].

**Figure 6.**
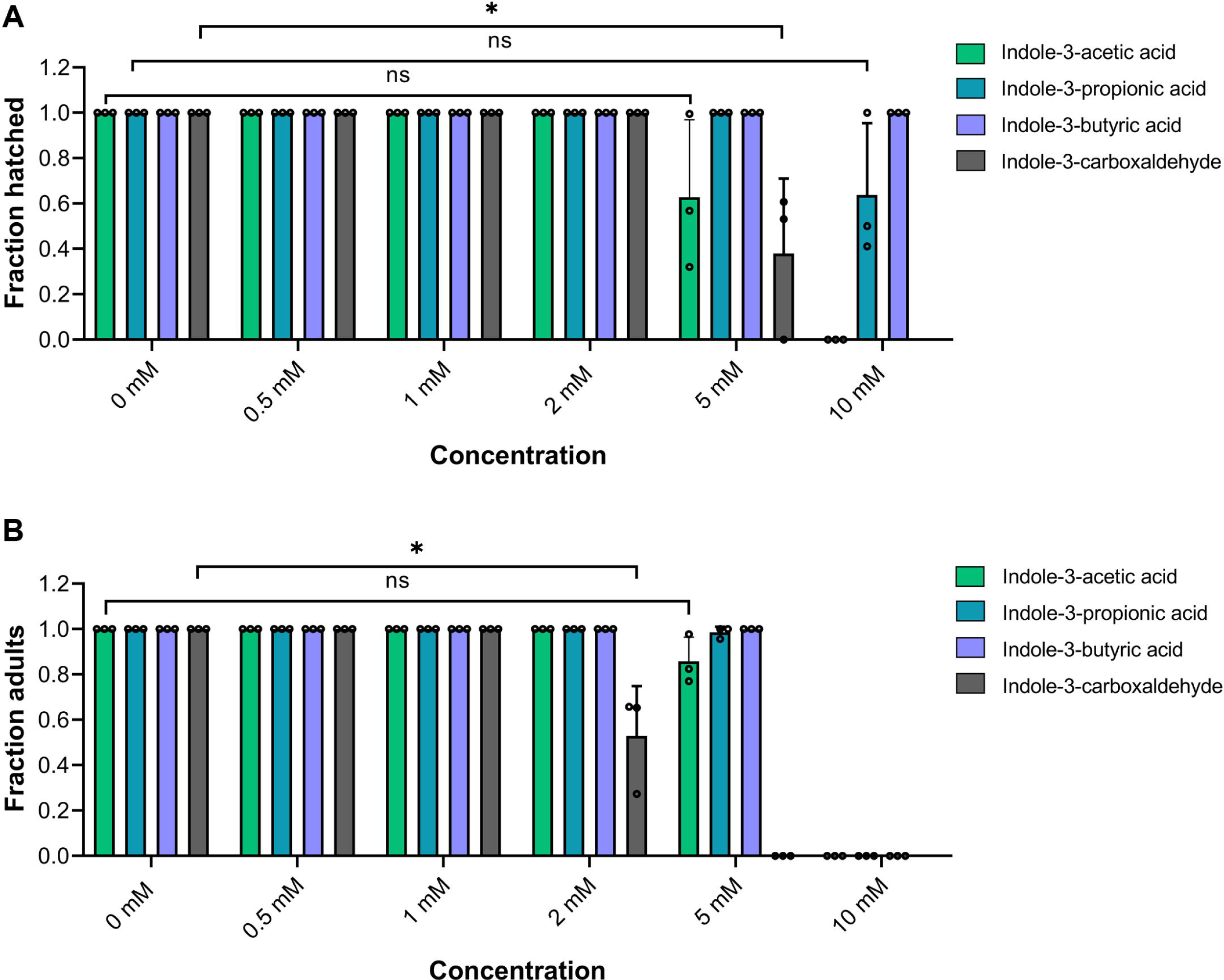
Indole derivatives cause toxicity in *C. elegans*. (A) Quantification of hatched eggs of N2 worms on various concentrations of indole-3-acetic acid, indole-3-propionic acid, indole-3-butyric acid, and indole-3-carboxaldehyde on *E. coli* OP50 diet after 24 hours of hatching at 20°C. The 0 mM control is the same for all the derivatives, as the experiments were done together. \**P* < 0.05 and n.s., nonsignificant via t-test (*n >*50 eggs for each condition from three independent biological replicates). (B) Quantification of adult fraction of N2 worms on various concentrations of indole-3-acetic acid, indole-3-propionic acid, indole-3-butyric acid, and indole-3-carboxaldehyde on *E. coli* OP50 diet after 96 hours of hatching at 20°C. The 0 mM control is the same for all the derivatives, as the experiments were done together. \**P* < 0.05 and n.s., nonsignificant via t-test (*n >*50 worms for each condition from three independent biological replicates). Indole-3- carboxaldehyde precipitates above 5 mM concentration in the NGM. Therefore, there is no 10 mM data for indole-3-carboxaldehyde in panels A and B.

### Indole does not cause toxicity via oxidative stress

Because genes induced by tryptophan and indole exposure were enriched for monooxygenase and oxidoreductase activities (Fig. 1E, F) [31], functions commonly associated with oxidative stress [32], we next investigated whether indole exerts its toxic effects through oxidative stress. Supplementation with the antioxidant N-acetylcysteine failed to alleviate tryptophan-induced toxicity (Fig. 7A), suggesting that oxidative stress is unlikely to be the primary cause. To further examine this possibility, we assessed the sensitivity of *C. elegans* mutants defective in oxidative stress responses (*skn-1*, *sek-1*, and *hlh-30*) and in the mitochondrial unfolded protein response (*atfs-1*) to indole exposure. While *skn-1(zj15)* and *atfs-1(gk3094)* mutants exhibited a modest baseline reduction in egg hatching (∼20%) on control plates, indole treatment did not further increase their sensitivity relative to wild-type N2 animals (Fig. 7B). Similarly, *hlh-30(tm1978)* mutants displayed no enhanced susceptibility to indole toxicity. Unexpectedly, *sek-1(km4)* and *atfs-1(gk3094)* mutants were more resistant to indole, producing a significantly higher proportion of hatched eggs at 1 mM indole compared to N2 worms (Fig. 7B). Together, these findings indicated that indole toxicity in *C. elegans* is not mediated by oxidative stress.

**Figure 7.**
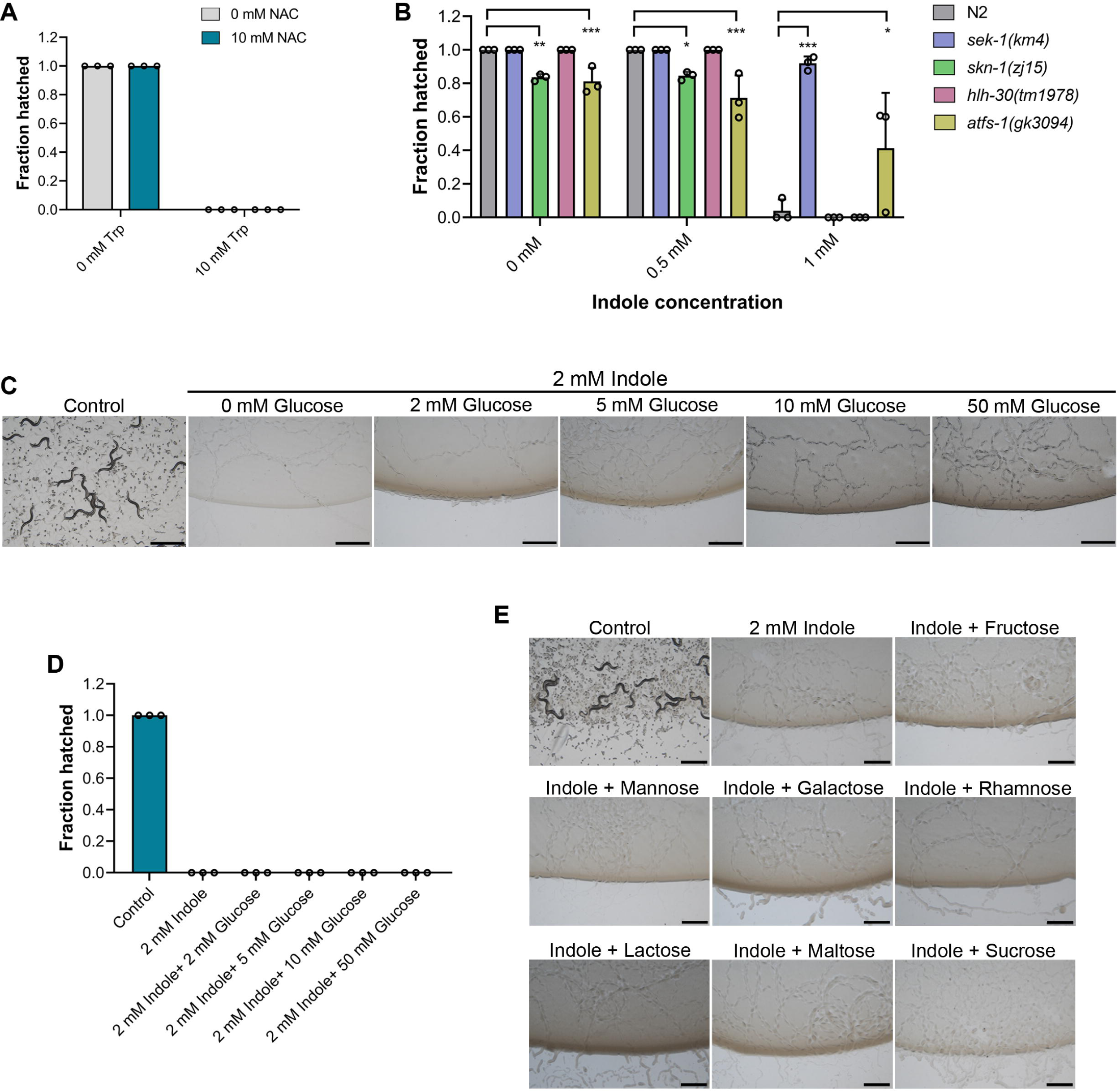
Indole does not cause toxicity by oxidative stress, and sugars do not rescue indole toxicity. (A) Quantification of hatched eggs of N2 worms on 10 mM tryptophan and 10 mM tryptophan supplemented with 10 mM N-acetylcysteine (NAC) on *E. coli* OP50 after 24 hours of hatching at 20°C (*n >*200 eggs for each condition from three independent biological replicates). (B) Quantification of hatched eggs of N2, *sek-1(km4)*, *skn-1(zj15)*, *hlh-30(tm1978)*, and *atfs- 1(gk3094)* worms on 0 mM, 0.5 mM, and 1 mM indole on *E. coli* OP50 diet for 24 hours at 20°C. \**P* < 0.05, \*\**P* < 0.01, and \*\*\**P* < 0.001 via ordinary one-way ANOVA followed by Dunnett’s multiple comparisons test (*n >*50 eggs for each condition from three independent biological replicates). (C) Representative images of N2 worms grown on *E. coli* OP50 diet for 96 hours at 20°C containing 0 mM indole (control), 2 mM indole, and 2 mM indole supplemented with increasing concentration of glucose. Scale bar = 1 mm. (D) Quantification of hatched eggs of N2 worms grown on *E. coli* OP50 diet for 24 hours at 20°C containing 0 mM indole (control), 2 mM indole, and 2 mM indole supplemented with different concentrations of glucose (*n >*150 eggs for each condition from three independent biological replicates). (E) Representative images of N2 worms grown on *E. coli* OP50 diet for 96 hours at 20°C containing 0 mM indole (control), 2 mM indole, and 2 mM indole supplemented with 50 mM of different sugars. Scale bar = 1 mm.

### Sugars do not alleviate indole-induced toxicity in *C. elegans*

We next sought to understand the mechanism by which glucose and other sugars suppress tryptophan toxicity. Transcriptomic profiling (Table S1) revealed that tryptophan exposure leads to the upregulation of numerous UGT genes. Notably, the transcriptional response to tryptophan closely resembles that elicited by indole exposure, as reported previously [31], suggesting that UGT induction may constitute a host defense mechanism against indole-mediated toxicity. UGT enzymes play a central role in phase II xenobiotic detoxification by conjugating glucose moieties to xenobiotic substrates, thereby increasing their solubility and facilitating excretion. Consistent with this, *C. elegans* has been shown to glycosylate indoles to reduce their toxicity [33].

We hypothesized that glucose supplementation may enhance UGT-mediated detoxification by increasing the availability of glucose substrates for conjugation. Under this model, the glycosylation of indoles could be responsible for the observed rescue of tryptophan toxicity when glucose is added. To test this hypothesis, we supplemented indole-containing plates with glucose and assessed egg hatching. Unexpectedly, even 50 mM glucose failed to promote hatching on plates containing 2 mM indole (Fig. 7C, D), indicating that glucose supplementation does not alleviate indole toxicity. To further evaluate this, we supplemented indole plates with other sugars that were tested for mitigating tryptophan toxicity. However, none of these sugars ameliorated the toxic effects of indole (Fig. 7E).

These results suggested that sugar-mediated rescue of tryptophan toxicity does not occur via enhanced UGT-mediated glycosylation of indole, and that sugars do not protect *C. elegans* from indole toxicity.

### Sugars alleviate tryptophan toxicity by suppressing bacterial indole production

We observed that several sugars alleviate the toxic effects of tryptophan but not those of indole. This suggests that these sugars may act upstream of indole production, potentially preventing tryptophan from being converted into its toxic derivative. We therefore asked whether sugar supplementation affects bacterial indole production. Previous studies have shown that glucose represses the expression of TnaA and can also inhibit its activity [34,35]. Based on these findings, we hypothesized that supplementation of sugars might reduce indole production even in the presence of excess tryptophan by repressing TnaA.

To test this, we quantified indole production by bacteria using Kovac’s reagent. As expected, *E. coli* cultured in LB medium produced indole under basal conditions, and indole levels were further elevated upon tryptophan supplementation (Fig. 8A). Notably, glucose supplementation significantly reduced indole production in cultures exposed to high concentrations of tryptophan (Fig. 8A). We extended this analysis to other monosaccharides and disaccharides. Strikingly, only those sugars that rescued tryptophan toxicity also reduced indole production. In contrast, sucrose and maltose, both ineffective at rescuing tryptophan toxicity, did not significantly lower indole levels compared to the tryptophan-only control (Fig. 8A). Furthermore, the sugars that alleviated tryptophan toxicity also diminished basal indole production in the absence of exogenous tryptophan (Fig. 8B).

**Figure 8.**
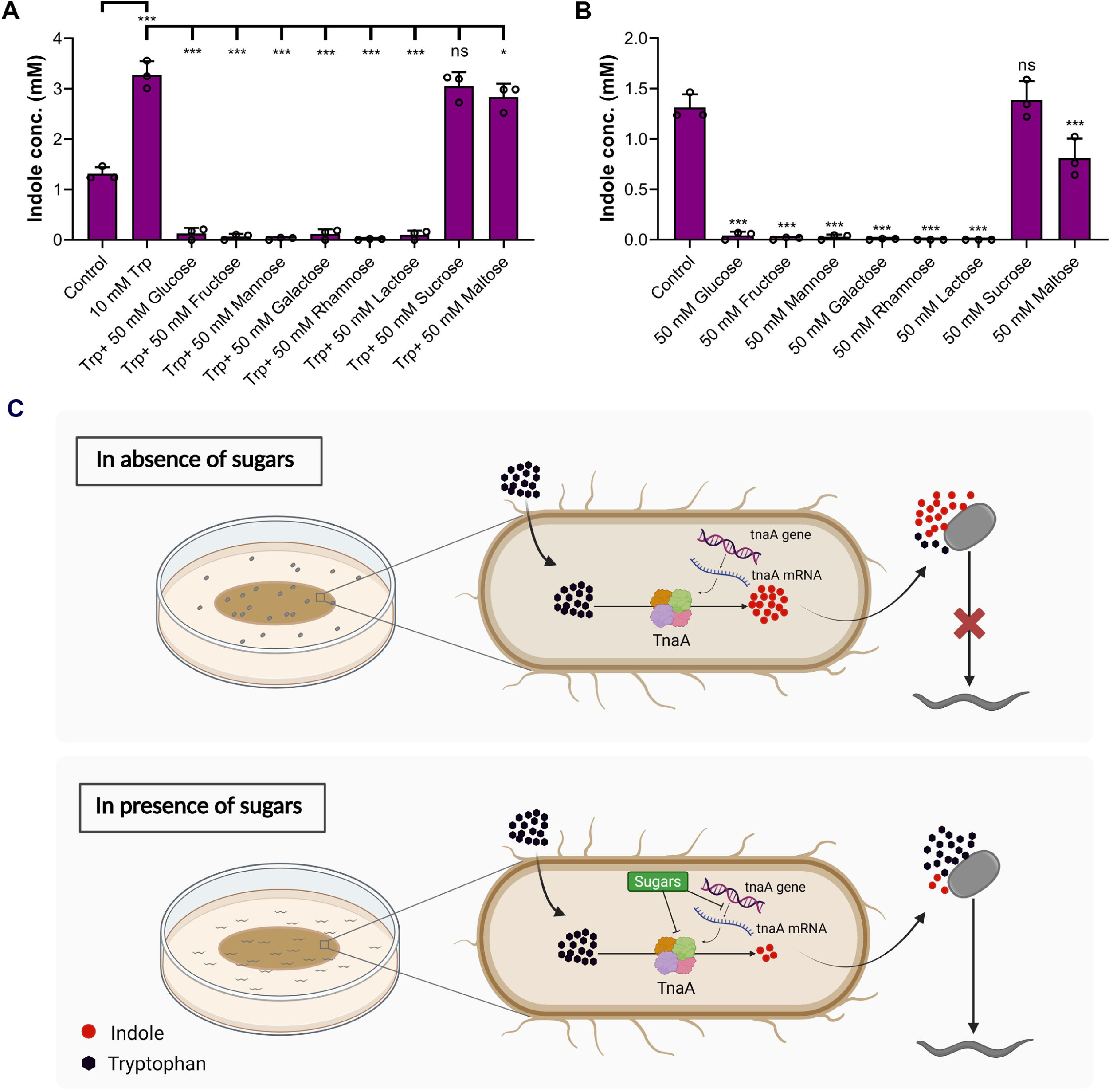
Sugars alleviate tryptophan toxicity by suppressing bacterial indole production. (A) Quantification of indole produced by *E. coli* OP50 cultured without any supplement (control), with 10 mM tryptophan (Trp), and with 10 mM Trp plus 50 mM of various sugars. \**P* < 0.05, \*\*\**P* < 0.001, and n.s., nonsignificant via ordinary one-way ANOVA followed by Dunnett’s multiple comparisons test (*n =* 3 biological replicates). (B) Quantification of indole produced by *E. coli* OP50 cultured without any supplement (control) and with 50 mM of various sugars. \*\*\**P* < 0.001 and n.s., nonsignificant via ordinary one-way ANOVA followed by Dunnett’s multiple comparisons test (*n =* 3 biological replicates). The control data in (A) and (B) are the same. (C) Model showing that the bacterial metabolism of tryptophan causes toxicity in *C elegans*. *E. coli* takes up tryptophan and catabolizes it to indole via tryptophanase (TnaA). Indole can freely diffuse through lipid membranes and prevents hatching of worm eggs at high concentrations. The addition of sugars decreases indole production from tryptophan by repressing the expression of *tnaA* in *E. coli*, resulting in the hatching and development of *C. elegans* in the presence of excessive tryptophan.

Collectively, these findings support a model in which *E. coli* converts tryptophan to indole, which is then taken up by *C. elegans*. At high concentrations, indole is toxic and impairs egg hatching. Various sugars, including multiple monosaccharides and the disaccharide lactose, inhibit indole biosynthesis in *E. coli*, thereby preventing the downstream toxic effects of tryptophan in *C. elegans* (Fig. 8C).

## Discussion

Our study demonstrates that bacterial metabolism of tryptophan into indole can profoundly impact host physiology and contribute to toxicity. Over 85 bacterial species, including both Gram-positive and Gram-negative, have been reported to express functional tryptophanase and convert tryptophan to indole [24]. Many of these species reside in the human gut, where they efficiently produce indole and its derivatives [36]. Human fecal samples have been reported to contain approximately 10 mM tryptophan [37], providing an abundant substrate for bacterial indole production. Indeed, indole is the most abundant tryptophan-derived metabolite detected in human feces and mouse cecal contents, accounting for 50-75% of total tryptophan catabolites, with concentrations of 2.6-2.7 mM [38,39]. The indole concentrations could increase further during enhanced tryptophan uptake, as indole concentrations as high as 6.6 mM have been observed in human faeces [38]. Therefore, indoles produced by gut microbiota might lead to toxicity under specific circumstances.

We observed that tryptophan induced toxicity in *C. elegans* not only during egg hatching but also in adults, causing paralysis. Interestingly, previous studies have reported that elevated tryptophan levels, achieved either by supplementation or inhibition of its catabolism, extend *C. elegans* lifespan [8–10]. These apparent discrepancies are likely attributable to differences in experimental design. Inhibition of tryptophan catabolism extends lifespan presumably by elevating internal tryptophan levels within the host, without increasing the pool available to gut bacteria for indole production. Moreover, studies reporting lifespan extension upon tryptophan supplementation employed heat-killed bacteria as a food source, thereby eliminating bacterial tryptophan metabolism [9]. Together, these observations suggest that tryptophan itself is beneficial to *C. elegans*, whereas its bacterial conversion to indole underlies the observed toxic effects.

Indole and its derivatives exhibit pleiotropic effects on host biology, ranging from beneficial to detrimental, depending on the context [40–44]. At moderate concentrations, indole can promote host health. For example, 250 µM indole has been shown to extend healthspan in *C. elegans*, *Drosophila*, and mice via conserved molecular pathways [45,46]. Indole also supports intestinal homeostasis by enhancing epithelial barrier function, modulating immune responses, and suppressing the colonization of pathogenic bacteria [47–50]. Furthermore, indole and its derivatives have been implicated in ameliorating inflammatory bowel disease, hemorrhagic colitis, colorectal cancer, and type 2 diabetes mellitus [29].

However, at elevated concentrations or in the context of gut dysbiosis, indole and its metabolites can become deleterious. In the liver, indole is converted to indoxyl sulfate, a known uremic toxin linked to systemic toxicity. Indoxyl sulfate has been implicated in the pathogenesis of cardiovascular and chronic kidney diseases [51–53], and is capable of inducing neuroinflammation and neuronal damage [54,55]. Additionally, indole exacerbates portal hypertension in a rat model of liver cirrhosis [56] and can promote the development of collagen-induced arthritis in mice via microbiota-dependent mechanisms [44]. In *C. elegans*, bacterial expression of the tryptophanase gene has been linked to bacterial virulence [57], and elevated indole levels have been shown to reduce lifespan [31,58]. These observations underscore that the pathological consequences of indole are context-dependent, influenced by its concentration and specific derivatives.

A key question emerging from our study concerns the mechanisms underlying indole toxicity. Indole is known to freely diffuse across lipid bilayers [30] and can disrupt membrane potential [59]. In bacteria, such perturbation of membrane potential inhibits cell division [59]. Because indole also inhibits egg hatching in *C. elegans*, it may similarly interfere with eukaryotic cell division. Consistent with this notion, indole inhibits the growth of the eukaryotic parasite *Cryptosporidium parvum* by reducing membrane potential [60]. Thus, as in prokaryotes, indole may inhibit eukaryotic cell division by altering membrane potential and bioenergetics. Although indole has been reported to induce oxidative stress [61], our findings indicate that oxidative stress is unlikely to account for its toxicity. Beyond membrane potential disruption, indole also impairs mitochondrial function and decreases total cellular ATP levels in eukaryotic cells [60,62]. Further studies are therefore warranted to elucidate the precise mechanisms by which indole and its derivatives exert cytotoxic effects in eukaryotic systems.

Our findings reveal that several monosaccharides, as well as the disaccharide lactose, inhibit indole production in *E. coli*. Previous studies have shown that glucose and arabinose suppress indole production by repressing TnaA expression [34,35]. Although catabolite repression via cyclic AMP (cAMP) was initially proposed as the underlying mechanism [63], subsequent studies have shown that tryptophanase regulation also involves cAMP-independent pathways [35]. Tryptophanase activity is regulated through complex and multilayered networks [35,63–65], suggesting that indole production may offer evolutionary advantages to bacteria [66]. Indeed, indole acts as a conserved intercellular signaling molecule that modulates bacterial community behavior and composition [24,67]. It regulates diverse aspects of bacterial physiology, including spore formation, drug resistance, plasmid stability, virulence, and biofilm formation [24]. Consequently, protein- and tryptophan-rich diets may selectively enrich indole-producing bacteria in the gut microbiota, potentially predisposing the host to dysbiosis and associated pathophysiological outcomes.

Our results suggest that indole production by gut microbiota can be modulated through dietary interventions. Indeed, dietary components such as fiber, non-starch polysaccharides, and high-fat diets have been shown to reduce gut indole production, shifting tryptophan metabolism toward more beneficial pathways [68–71]. We identified additional sugars capable of suppressing indole synthesis in *E. coli* and possibly in other bacterial species. Future studies examining the interplay between dietary sugars and tryptophan metabolites in human gut contents, across varying diets, ages, and disease states, could yield valuable insights into microbiota metabolic states and their influence on host physiology.

## Materials and methods

### Bacterial strains

The following bacterial strains were used in this study: *Escherichia coli* OP50, *E. coli* BW25113, *E. coli* BW25113 *ΔtnaA* mutant, and *Pseudomonas aeruginosa* PA14 *ΔgacA* mutant. Bacterial cultures were grown in Luria-Bertani (LB) broth at 37°C. The *E. coli* BW25113 *ΔtnaA* mutant was from the *E. coli* Keio collection [72], and was cultured in LB broth supplemented with 25 µg/mL kanamycin. Because *P. aeruginosa* virulence might affect the hatching and development of *C. elegans*, we used the *gacA* mutant, which has drastically reduced virulence [73]. Nematode growth medium (NGM) plates were seeded with the respective bacterial cultures and incubated at room temperature for at least two days prior to use in experiments.

### *C. elegans* strains and growth conditions

*C. elegans* hermaphrodites were maintained at 20°C on NGM plates seeded with *E. coli* OP50 as described earlier [74,75]. The Bristol N2 strain was used as the wild-type control. The following strains were used in this study: RG3052 *tdo-2(ve552[LoxP + myo- 2p::GFP::unc-54 3’ UTR + rps-27p::neoR::unc-54 3’ UTR + LoxP])*, RB1190 *amx-2(ok1235)*, MT15434 *tph-1(mg280)*, KU4 *sek-1(km4)*, QV225 *skn-1(zj15)*, JIN1375 *hlh-30(tm1978)*, and VC3201 *atfs-1(gk3094)*. Some of the strains were obtained from the Caenorhabditis Genetics Center (University of Minnesota, Minneapolis, MN).

### Preparation of NGM plates with different supplements

The following supplements, indicated with their product numbers, were obtained from HiMedia BioSciences: L-tryptophan (#GRM067), D-(+)-glucose (#TC130), D-(-)-fructose (#GRM196), D-(+)-mannose (#RM104), L-(+)-rhamnose (#RM062), D-(+)-galactose (#GRM101), lactose (#GRM017), sucrose (#GRM134), D-(+)-maltose (#GRM3050), indole (#GRM824), indole-3-acetic acid (#RM384), indole-3-propionic acid (#PCT0831), indole-3- butyric acid (#RM385), indole-3-carboxaldehyde (#RM2874), sodium pyruvate (#GRM1181), ammonium chloride (#MB054), and N-acetylcysteine (#RM3142). Indole, indole-3-acetic acid, indole-3-propionic acid, indole-3-butyric acid, and indole-3- carboxaldehyde were dissolved in methanol, and control plates were supplemented with equivalent volumes of methanol for assays involving indole and its derivatives.

Tryptophan powder was weighed, suspended in water, and added directly to molten NGM. Stock solutions of 1 M were prepared in water for D-(+)-glucose, D-(-)-fructose, D- (+)-mannose, L-(+)-rhamnose, sucrose, and D-(+)-maltose. A 0.5 M aqueous stock was prepared for D-(+)-galactose, whereas lactose powder was added directly to the medium. Sodium pyruvate (1 M) and ammonium chloride (2 M) stocks were also prepared in water. Stock solutions (250 mM) of indole, indole-3-acetic acid, indole-3-propionic acid, and indole-3-butyric acid were prepared in methanol, and indole-3-carboxaldehyde was prepared as a 100 mM stock in methanol. N-acetylcysteine (0.5 M) was prepared in water, and its pH was adjusted to 6.1 before addition to the medium. All supplements were added to the NGM immediately before pouring plates to achieve the desired final concentrations.

### *C. elegans* egg hatching assays

A synchronized egg population was obtained by transferring 15-20 gravid adult hermaphrodites onto NGM plates and allowing them to lay eggs for 1.5-2 hours. For *sek- 1(km4)*, *skn-1(zj15)*, *hlh-30(tm1978)*, and *atfs-1(gk3094)* mutants, 25-30 gravid adults were transferred for egg laying for 2-2.5 hours. After that, the adults were removed, and the number of eggs laid was counted. The plates were then incubated at 20°C, and the number of hatched eggs/L1 larvae was quantified after 24 hours. Representative images of the plates were captured after 96 hours of incubation at 20°C. For each experimental condition, at least three independent biological replicates were performed.

### Paralysis assays

Approximately 50-60 gravid adult worms were transferred to NGM plates supplemented with varying concentrations of tryptophan, or with 10 mM tryptophan in combination with 50 mM glucose. NGM plates without any supplements served as controls. Worms were examined manually under a stereomicroscope at 30-minute intervals to assess paralysis. For each condition, at least three independent biological replicates were performed.

### RNA isolation

RNA isolation was performed as described previously [76,77]. Briefly, wild-type N2 animals were synchronized by egg laying. Approximately 50 gravid adult hermaphrodites were transferred to 9-cm NGM plates seeded with *E. coli* OP50 and allowed to lay eggs for 4 hours. After removing the adults, the eggs were incubated at 20°C for 72 hours to allow development into adult worms. The adult animals were then collected, washed with M9 buffer, and transferred to *E. coli* OP50-seeded plates containing the following three conditions: control plates without supplements, 2 mM tryptophan, and 2 mM tryptophan plus 50 mM glucose. Following a 6-hour incubation at 20°C, worms were collected, washed with M9 buffer, and snap-frozen in TRIzol reagent (Life Technologies, Carlsbad, CA). Total RNA was extracted using the RNeasy Plus Universal Kit (Qiagen, Netherlands). The RNA samples from three independent biological replicates were isolated for each condition.

### RNA sequencing and data analysis

Total RNA was isolated from N2 worms grown under three conditions: control, 2 mM tryptophan, and 2 mM tryptophan plus 50 mM glucose, using three biological replicates per condition, as described above. Library preparation and sequencing were conducted at Unipath Specialty Laboratory Ltd., India. The cDNA libraries were sequenced on the NovaSeq 6000 platform using 150-bp paired-end reads.

The RNA sequencing data were analyzed using the web platform Galaxy (https://usegalaxy.org/), following previously described workflows [78,79]. Briefly, paired-end reads were first trimmed using the Trimmomatic tool. The resulting reads obtained for each sample were mapped to the *C. elegans* reference genome (WS220) using the STAR aligner. Read counts per gene were quantified using the htseq-count tool. Differential gene expression analysis was then performed using DESeq2. Genes exhibiting at least a two-fold change and a *P*-value <0.01 were considered differentially expressed. Gene ontology enrichment analysis was carried out using the DAVID Bioinformatics Resource (https://david.ncifcrf.gov/tools.jsp). Venn diagrams were generated using the BioVenn web tool (https://www.biovenn.nl/), and heatmaps were created using the pheatmap package in R version 4.5.0 (R Statistical Software).

### Bacterial killing

Bacterial killing was performed as described previously [80]. Briefly, *E. coli* OP50 was cultured in 50 mL of LB broth at 37°C for 24 hours. The culture was then centrifuged at 3000 × *g* for 30 minutes. The supernatant was discarded, and the bacterial pellet was resuspended in 100 mL of autoclaved water containing 250 µg/mL kanamycin. The suspension was distributed equally into two 250-mL flasks and incubated with shaking at 37°C for an additional 24 hours. To confirm bacterial killing, 1 mL of the culture was aliquoted, washed three times with sterile water, and plated on LB agar. Plates were incubated at 37°C for 24 hours and checked for viable colonies. If no colonies were detected, the bacteria were considered effectively killed and stored at 4°C. Before use, the killed bacterial culture was washed three times with 100 mL of sterile water and concentrated 20-fold before seeding onto NGM plates.

### Indole measurement using Kovac’s reagent

To prepare samples for indole quantification, various sugars were added to LB broth at a final concentration of 50 mM. Each sugar was added to 10 mL of LB broth in 50 mL tubes, with or without added tryptophan. After that, 20 µL of the same primary culture of *E. coli* OP50 was added into each tube, which were then incubated at 37°C for 12 hours. The optical density at 600 nm (OD_600_) was measured for all cultures, and the cultures were adjusted to the same OD_600_ prior to indole quantification. As a control, 10 mL of LB broth containing *E. coli* OP50 without added sugar or tryptophan was similarly cultured. Indole standards (0, 0.1, 0.5, 1, 2, 5, and 10 mM) were prepared in LB broth and processed in parallel with the samples to generate a calibration curve.

Indole levels were measured as described previously [81], with minor modifications.

Briefly, 1 mL of each culture was transferred to a 1.5 mL microcentrifuge tube and centrifuged at 11,300 × *g* for 20 seconds. Then, 250 µL of the supernatant was mixed with 250 µL of 20% (w/v) trichloroacetic acid (Sigma-Aldrich #T6399) and incubated on ice for 15 minutes. The mixture was centrifuged at 13,000 × *g* for 10 minutes to remove precipitated proteins. Subsequently, 450 µL of the resulting supernatant was combined with 500 µL of Kovac’s reagent (Sigma-Aldrich #67309) and vortexed. The samples were incubated at room temperature for 2 minutes to allow phase separation. Then, 200 µL of the upper (colored) layer was transferred to 800 µL of ethanol, and absorbance was measured at 540 nm. Indole concentrations were calculated using a standard curve generated on the same day.

### Quantification and statistical analysis

The statistical analysis was performed with Prism 8 (GraphPad). All error bars represent the mean ± standard deviation. The unpaired, two-tailed, two-sample *t*-test was employed when needed, and statistical significance was determined when *P* < 0.05. For more than two samples, ordinary one-way ANOVA followed by Dunnett’s multiple comparisons test was used. In the figures, asterisks (*) denote statistical significance as follows: \**P* < 0.05; \*\**P* < 0.01; and \*\*\**P* < 0.001, as compared with the appropriate controls. All experiments were performed at least three times.

## Acknowledgments

We thank the Caenorhabditis Genetics Center [funded by the NIH Office of Research Infrastructure Programs (P40 OD010440)] for providing the strains used in this study. We thank Ritika Bassi (IISER Mohali) for help with the generation of the heatmap.

## Author Contributions

S.G., Subodh, and J.S. conceived and designed the experiments. S.G. and Subodh performed the experiments. S.G. and J.S. analyzed the data and wrote the paper.

## Data availability

The RNA sequencing data for N2 worms exposed to tryptophan alone and tryptophan plus glucose, along with the control, have been submitted to the public repository, the Sequence Read Archive, with BioProject ID PRJNA1271509. All data generated or analyzed during this study are included in the manuscript and supporting files.

## Abbreviations

ATP,: Adenosine triphosphate;
cAMP,: cyclic AMP;
GO,: gene ontology;
LB,: Luria-Bertani;
NAC,: N-acetylcysteine;
NGM,: nematode growth medium;
TnaA,: tryptophanase;
Trp,: tryptophan;
UGT,: UDP-glucuronosyltransferase

## Supplementary Tables

**Table S1.** Differentially-regulated genes in adult N2 worms exposed to 2 mM tryptophan for 6 hours. A separate Excel file is provided.

**Table S2.** Differentially-regulated genes in adult N2 worms exposed to 2 mM tryptophan plus 50 mM glucose for 6 hours. A separate Excel file is provided.

**Table S3.** Comparison of genes upregulated in N2 worms upon exposure to tryptophan alone and tryptophan plus glucose. A separate Excel file is provided.

**Table S4.** Comparison of genes downregulated in N2 worms upon exposure to tryptophan alone and tryptophan plus glucose. A separate Excel file is provided.

